# Interplay Between KRAS and LZTR1 Protein Turnover, Controlled by CUL3/LZTR1 E3 Ubiquitin Ligase, is Disrupted by KRAS Mutations

**DOI:** 10.1101/2021.11.23.469679

**Authors:** Andreas Damianou, Zhu Liang, Frederik Lassen, George Vere, Svenja Hester, Philip D Charles, Adan Pinto-Fernandez, Alberto Santos-Delgado, Roman Fischer, Benedikt M Kessler

**Author notes:** Corresponding authors: Andreas Damianou Benedikt Kessler TDI, CAMS-COI, CMD, Nuffield Department of Medicine University of Oxford, Roosevelt Drive Oxford OX3 7FZ, UK. Equal contribution (Andreas Damianou, Zhu Liang).

## Abstract

KRAS is a proto-oncogene encoding a small GTPase. Mutations contribute up to 30% of human solid tumours including lung adenocarcinoma, pancreatic and colorectal carcinomas. Most KRAS activating mutations interfere with GTP hydrolysis, essential for its role as a molecular switch, leading to alterations in their molecular environment and oncogenic signalling. Here, APEX-2 proximity labelling was used to profile the molecular environment of wild type and G12D, G13D and Q61H activating mutants of KRAS under both, starvation and stimulation conditions. We demonstrate by quantitative proteomics the presence of known interactors of KRAS including a-RAF and LZTR1, which varied in abundance with wildtype and KRAS mutants. Notably, the KRAS mutations G12D, G13D and Q61H abrogate association with LZTR1. Wildtype KRAS and LZTR1, as part of the CUL3 ubiquitin E3 ligase complex, affect each other’s protein stability, revealing a direct feedback loop mechanism. KRAS mutations disconnect this regulatory circuit, thereby contributing to oncogenesis.

## INTRODUCTION

Kirsten rat sarcoma virus (KRAS) is a signal transducing proto-oncogene that belongs to the superfamily of small GTPases **(Gimple and Wang, 2019)**. KRAS and other RAS family proteins are molecular switches that alternate between an active and inactive conformational state, determined by its binding to GTP or GDP respectively. Upon environmental stimuli, such as GF mediated signalling, guanine nucleotide-exchange factors (GEF’s), including RASGEF1A and RASGRF2 are recruited to facilitate the exchange of GDP to GTP, leading to activation of KRAS **(Buday and Downward, 1993) (Vigil et al., 2010).** Active KRAS promotes signal transduction through effectors such as RAF **(Moodie et al., 1993)**, TIAM1 **(Lambert et al., 2002)** and PI3K **(Castellano and Downward, 2010)** and persists until GTP is hydrolysed to GDP, which is catalysed by KRAS-associated GTPase-activating proteins (GAPs).

A balanced equilibrium exists between active and inactive RAS, including KRAS, HRAS and NRAS. In healthy mammalian cells, this can be disrupted by activating point mutations in KRAS, leading to neoplastic properties **(Prior et al., 2012)**. These mutations activate chronic KRAS signalling, either by impairing GTPase activity and/or the ability of KRAS to interact with negative regulators **(Liu et al., 2019)**. Subsequent constitutive activation of pro-neoplastic signalling leads to uncontrolled cell division and malignant transformation **(Haigis, 2017)**.

Despite these detailed insights into KRAS as a GTPase, its turnover and mutation driven effects on KRAS interactors remain incompletely understood. Proximity labelling techniques, such as BioID, TurboID and ascorbate peroxidase 2 (APEX-2), developed recently to capture of protein-protein interactions as well as the exploration of different molecular and cellular environmental ‘proximitome’ **(Trinkle-Mulcahy, 2019)**, were applied to wildtype KRAS. For instance, a BioID study investigated the interactome of KRAS, unravelling mTORC2 as a direct KRAS effector **(Kovalski et al., 2019)** as well as LZTR1, a ubiquitin E3 ligase adaptor **(Bigenzahn et al., 2018)**. LZTR1, as part of the CUL3 E3 ligase complex, was demonstrated to ubiquitinate HRAS at position K170, which consequently alters its cellular localisation by attenuating the interaction of HRAS with the membrane **(Steklov et al., 2018)**. Additionally, LZTR1 inactivation can lead to a decrease in KRAS ubiquitination, which promotes localisation of KRAS to the plasma membrane **(Bigenzahn et al., 2018)**. Both reports suggest that LZTR1 affects KRAS function via a non-degradative ubiquitination mechanism. In addition, LZTR1 was proposed also to function also as a “Ras killer protein” by inducing polyubiquitination and degradation of endogenous RAS via the ubiquitin-proteasome pathway **(Abe et al., 2020)**. An additional level of regulation was revealed by KRAS mono-ubiquitylation at K104, which restores GEF-mediated nucleotide exchange **(Yin et al., 2020).**

Whereas previous studies have predominantly focused on wildtype KRAS, we wondered how oncogenic mutations may affect dynamic proximal molecular interactions. To this end, we applied APEX-2, a rapid proximity labelling approach, to compare interactomes of wildtype KRAS with three mutants G12D, G13D and Q61H upon starvation and/or acute stimulation by exposure to foetal calf serum (FCS) to reveal dynamic interactions of active and inactive forms. Quantitative mass spectrometry analysis confirmed known KRAS interactors such as RAF as well as LZTR1. Strikingly, the wildtype KRAS proximal proteome was altered strongly between the starvation (GDP-bound form) versus the FCS induced (GTP-bound), a trait that was less pronounced with mutants. When comparing the proximal proteome of wildtype KRAS with three different constitutively active mutants, LZTR1 was one of the most predominant proteins preferentially interacting with wildtype but not the mutants. Biochemical studies uncovered that wildtype KRAS and LZTR1, as part of the CUL3 ubiquitin E3 ligase complex, affect each other’s protein stability, revealing a direct feedback loop mechanism. KRAS activation mutations appear to disconnect this regulatory circuit, reflecting a possible contribution to tumorigenesis.

## MATERIALS AND METHODS

### Cell lines

Flp-In™ T-REx™ 293 Cell Line (parental line; R78007, ThermoFisher UK) were cultured in D-MEM™ (high glucose, Gibco #12440-53) supplemented with 10% FBS (Gibco #10500-64), 2 mM L-glutamine (G7513, Sigma-Aldrich), 100 µg/mL Zeocin™ (InvivoGen, ant-zn-1) 15 µg/mL blasticidin (InvivoGen, ant-bl-1) at 37 °C in a humidified 5% CO atmosphere. The stable transfected Flp-In™ T-REx™ 293 Cell Lines (KRas WT-APEX2, KRas G12D -APEX2, KRas G13D -APEX2, KRas Q61H –APEX2, KRas WT-GFP, KRas G12D - GFP, KRas G13D -GFP, KRas Q61H –GFP, GFP only) were cultured in the same medium and selected by adding 100 μg/ml of hygromycin B (InvivoGen, ant-hg-1) to the media. For the induction of the stable transfected Flp-In™ T-REx™ 293 Cell Lines Cells 1μg/mL tetracycline was added to the media.

### Plasmids

Q5® High-Fidelity DNA Polymerase (NEB, M0491S) was used to perform PCR reactions following the manufacturer’s instructions. APEX2 plasmid was donated from Pedro Carvalho’s lab (pcDNA3 APEX2- NES). APEX2 ORF was then PCR amplified and inserted into pcDNA™5/FRT Vector (Invitrogen™) using NEBuilder® HiFi DNA Assembly (NEB, E5520S) following the manufacturer’s protocol. KRAS ORF was initially codon optimised and synthesized from Eurofins Scientific. Then KRAS ORF was PCR amplified and cloned into the pcDNA™5/FRT Vector APEX-2 plasmid using NEBuilder® HiFi DNA Assembly following protocol. Different KRAS mutants were generated using Q5® Site-Directed Mutagenesis Kit (NEB, E0554S). The successful clones were confirmed using DNA sequencing.

A pcDNA5/FRT/TO GFP (ITEM 19444) was purchased from Addgene. PCR was used to amplify all the different KRAS (WT, G12D, G13D, Q61H) ORF as well as the pcDNA5/FRT/TO GFP. NEBuilder® HiFi DNA Assembly was used to generate the GFP tagged plasmids. The successful plasmids were confirmed using DNA sequencing.

pcDNA3.1+ Flag-Kras-G12C was a kind gift from Dr Vikram Rao (Pfizer). WT, G12D, G13D and Q61H KRAS were generated using Q5® Site-Directed Mutagenesis Kit. The successful clones were confirmed using DNA sequencing. LZTR1 ORF and the pCI-backbone (Addgene, 41552) were PCR amplified and cloned using NEBuilder® HiFi DNA Assembly.

### Immunoblotting

5×10^6^ HEK293T or HEK293 FRT were washed with ice-cold PBS and lysed with RIPA buffer containing protease and phosphatase inhibitors. For immunoblotting, 25 μg of protein was then fractionated on Tris–glycine SDS-PAGE gradient (4–15% acrylamide) gels (BioRad; #3450123), transferred onto nitrocellulose membranes (Millipore), and detected with the indicated antibodies using a LI-COR detection system. Primary antibody used in this study: DYKDDDDK Tag Antibody (Flag, Cell Signaling Technology, 2368), A-Raf Antibody (Cell Signaling Technology, 4432), LZTR1 (Santa Cruz Biotechnology, sc-390166), p44/42 MAPK (Erk1/2) (Cell Signaling Technology, 9107), Phospho-p44/42 MAPK (Erk1/2) (Thr202/Tyr204) (Cell Signaling Technology, 4370) GAPDH (Sigma-Aldrich, G8795).

### Generation of Stable cell lines

Flp-In™ T-REx™ 293 Cell Line (Invitrogen™) was transfected using Lipofectamine LTX following the manufacturer’s protocol. 300 μg/ml of hygromycin B was added the next day and cells were incubated for 2 weeks while medium was changed every 4 days. Finally, the successful population of cells was further used for the described experiments.

### Proximity Labelling

5×10^6^ HEK293 FRT were induced with 1 μg/mL tetracycline overnight. The next day cells were incubated with 500 μM of Biotinyl tyramide for 30 minutes. A final concentration of 1 mM H_2_ 0_2_ was added for 40 seconds. The cells were then washed 2 times with quencher buffer (10 mM sodium ascorbate, 10 mM sodium azide, 5 mM Trolox), 2 times with ice-cold PBS and finally with quencher buffer for a last time. The cells were lysed with RIPA buffer containing 1 mM PMSF, 5 mM Trolox, 10 mM Sodium Azide, 10 mM sodium ascorbate, protease and phosphatase inhibitors.

### Transient Transfection

For DNA plasmids: 5×10^6^ HEK293T cells were grown in 6-well plates and Lipofectamine™ 3000 Transfection Reagent (L3000001, ThermoFisher UK) was used to transfect the cells following the manufacturer’s instructions. The concentration of plasmids used was indicated in each experiment.

For siRNA transfection experiments: HEK293 cells were grown in 6-well plates and RNAimax transfection reagent (Invitrogen #13778-150) was used following the manufacturer’s protocol. A final concentration of 10 nM of the following siRNAs were used: LZTR1 siRNA on-TARGET plus SMARTpool (Dharmacon, L-012318-00-0005) and CUL3 siRNA on-TARGET plus SMARTpool (Dharmacon, L-010224- 00-0005).

### Cycloheximide chase experiment

5×10^6^ HEK293T and HEK293 FRT were transfected with the indicated plasmids for 24 hours. A fresh medium was added for an additional 24 hours and then 50 µg/mL cycloheximide (CHX) was added. Western blot was used to evaluate different expression of proteins.

### RAS activation kit

5×10^6^ HEK293 FRT were incubated with or without FCS for the indicated time. Cells were then washed with ice-cold PBS and lysed with the designated lysis buffer provided by the active Ras detection kit. Subsequent steps were performed as per the manufacturer’s protocol of the active Ras detection kit (Cell Signaling Technology, 8821).

### Live Cell Imaging

10,000 HEK293 FRT were seeded in 96 well plates (M33089, ThermoFisher UK) and induced with tetracycline for 24 hours. The next day cells were transfected with LZTR1 siRNA or Control siRNA. 72 hours later we removed the medium. Hoechst 33342 (H3570, ThermoFisher UK) was diluted in PBS in 1:2000 dilution and added to the cells. The cells were then incubated for 10 minutes in the incubator. Cells were imaged using the Opera Phenix microscope. Images were further analysed in Columbus Image Data Storage and Analysis System. The media Cell fluorescent per mean fluorescent per well was then determined for each condition. Finally, the siRNA control was used to normalize the data. The experiment was performed in biological triplicate, and the graph was generated using Prism software (GraphPad Prism 9.2.0).

### Sucrose gradient fractionation

HEK293 FRT cells were homogenized in isotonic buffer (0.25 M sucrose, 10 mM Tris-HCl, pH 7,5, 10 mM KCl, 1.5 mM MgCl2, and protease inhibitor cocktail). After homogenization, whole cell lysates were centrifuged at 10,000 g for 10 min to obtain the crude membranes. P10 Crude membranes whereas washed once with the same isotonic buffer and resuspended in the same buffer. 60% and 20% sucrose solutions were prepared in low salt buffer (10 mM Tris-HCl, pH 7.5, 10 mM KCl, 1.5 mM MgCl_2_) and a continuous sucrose gradient was generated using BIOCOMP Model 108 GRAGIENT MASTER according to manufacturer’s guidance. P10 Crude membranes was then loaded on top of the gradient and centrifuged at 170,000 g for 2 hrs. 12 fractions (500 μl1 mL per fraction) were collected from top to bottom.

### Membrane-cytoplasmic cellular fractionation

Fresh HEK293 FRT cell pellets were suspended in digitonin lysis buffer (50 mM HEPES pH 7.5, 10 mg/mL digitonin, 150 mM NaCl) and incubated for 30 minutes in 4 °C under spinning. The sample was spun at 6000xg for 5 min in 4 °C. The supernatant (cytoplasmic fraction) was moved to a new tube. The pellet was re-suspended in 0.3 % NP-40, 50 mM HEPES pH 7.5 and 150 mM NaCl and incubated on ice for 5 min. The samples were centrifuged at 1500 g for 5 minutes. The supernatant (membrane fraction) was transfer to a new tube and stored at -20 °C until analysis.

### Streptavidin immunoprecipitation

For immunoprecipitation (IP) experiments, Dynabeads™ MyOne™ Streptavidin T1 for western blot 30 μL was used and for mass spectrometry 110 μL. Initially the beads were washed twice with RIPA buffer once with 1M KCL, 0.1 M Na_2_ CO_3_ and 2 M Urea in 10 mM Tris-HCL, pH 8 and finally twice with RIPA buffer again. For Western blot biotinylated proteins were eluted using 60 μL of 2X loading buffer supplemented with 2 mM biotin and 20 mM DTT by boiling (98 °C) for 10 minutes. For Mass spectrometry, an on bead-based digestion protocol was used as described below.

### On-beads digestion and mass spectrometry sample preparation

Immunoprecipitated protein samples were denatured in 8 M Urea in 100 mM triethylammonium bicarbonate buffer (TEAB) for 30 minutes at room temperature. Afterwards, the samples were reduced with 10mM tris(2-carboxyethyl)phosphine (TCEP) for additional 30 minutes at room temperature. 50 mM iodoacetamide was added to the sample for 30 minutes at room temperature to alkylate the proteins. Samples were then diluted down to 1.5 M Urea with 50 mM TEAB. Finally, 250 ng dissolved in 50 mM TEAB and incubated in 37 °C overnight. The next day samples were desalted using a C18 solid phase cartridge (Sep-Pak, Waters) following the manufacturer’s protocol. Purified peptide eluates were dried by vacuum centrifugation and re-suspended in buffer A (98 % MilliQ-H20, 2 % CH3CN and 0.1 % trifluoroacetic acid - TFA) and stored at -20 °C until analysis.

### Liquid chromatography-tandem mass spectrometry (LC-MS/MS) analysis

LC-MS/MS analysis was performed essentially as described previously **(Fischer and Kessler, 2015)**. In brief, resuspended peptide material was trapped on an Acclaim™PepMap™ 100 C18 HPLC Column (PepMapC18; 300 μm x 5 mm, 5 μm particle size, Thermo Fischer) using solvent A (98 % MilliQ-H20, 2 % CH3CN and 0.1 % TFA) at a pressure of 60 bar and separated on an Ultimate 3000UHPLC system (Thermo Fischer Scientific) coupled to a QExactive mass spectrometer (Thermo Fischer Scientific). The peptides were separated on an Easy Spray PepMap RSLC column (75 μm i.d. x 2 μm x50 mm, 100 Å, Thermo Fisher) and then electrosprayed directly into an QExactive mass spectrometer (Thermo Fischer Scientific) through an EASY-Spray nano-electrospray ion source (Thermo Fischer Scientific) using a linear gradient (length: 60 minutes, 5 % to 35 % solvent B (0.1 % formic acid in acetonitrile), flow rate: 250 nL/min). The raw data was acquired on the mass spectrometer in a data-dependent mode (DDA). Full scan MS spectra were acquired in the Orbitrap (scan range 380-1800 m/z resolution 70000, AGC target 3e6, maximum injection time 100 ms). After the MS scans, the 15 most intense peaks were selected for HCD fragmentation at 28 % of normalised collision energy. HCD spectra were also acquired in the Orbitrap (resolution 17500, AGC target 1e5, maximum injection time128 ms) with first fixed mass at 100 m/z.

### Proteomic data analysis

Raw mass spectrometry (MS) data was processed using MaxQuant software (v1.6.14.0) and searched against human protein sequences in UniProt/KB (version). For data analysis, Perseus (version 1.6.15.0) and Significance Analysis of INTeractome (SAINT) workflows **(Choi et al., 2012)** were applied. The MS data generated in this study has been submitted to the PRIDE public repository with the accession number PXD029725.

### Data analysis

We characterized differentially enriched proteins in G12, G13 and Q61H mutants versus wildtype during starvation and FCS. To do this, we first used Perseus (version 1.6.15.0) to perform a student’s t-test to evaluate Log Fold Changes (Log2 FC) between wildtype and mutants. For each condition, we then carried forward the differentially expressed proteins with a log2(mutant/WT FC) > 0.5 or <-0.5 and - log10 (p value) > 1.3. Proteins that fulfilled the above filtering criteria were considered to be potential mutant- or wildtype KRAS-specific interacting proteins. Gene ontology enrichment analysis of differential proximal proteins for three mutants were performed using the Metascape web-based tool (www.metascape.org) **(Zhou et al., 2019)**. To further investigate if KRAS mutants G12D, G13D and Q61H differ in downstream signalling pathways and known interactors, we compared the protein components in different pathways (MEK-RAF, mTOR, PI3K, RALGDS, RASSF, and TIAM-RAC) and known KRAS interacting partners enriched with different mutants and wildtype KRAS. Protein abundance was shown as Log2 (LFQ) value in the heatmap shown in **Fig 3E**.

We also investigated the overlap between our significantly enriched proteins (log2 [mutant/WT FC] > 0.5 or <-0.5 and - log10 [p value] > 0.8) and various gene sets including i) GO molecular function, ii) GO biological processes iii) GO cellular compartments and iv) MSigDB Hallmark genes and v) CORUM complexes **(Giurgiu et al., 2019)**. To do this, we used Genoppi **(Pintacuda et al., 2021)** to perform a one-tailed hypergeometric overlap test to assess the probability of finding a given gene set among our significantly enriched proteins. We performed two different analyses. First, we performed a “global analysis”, where we assess the overlap of gene sets in interactors compared to other genes in the genome. Second, we performed a “conditional analysis” where we assess the overlap of gene sets in interactors compared to other proteins found in the experiment.

### Software developed for data analytics

All analyses were performed with R version 4.0.2. Workflows and scripts have been compiled in a custom R-package that can be found in the following repository https://github.com/frhl/KRAS-methods.

## RESULTS

### KRAS-APEX-2 wildtype, G12D, G13D and Q61H mutants express, localise and function similarly to endogenous KRAS

To comprehensively characterize the proximal proteome of wildtype (WT) KRAS and of three of the most frequent oncogenic driver mutants (G12D, G13D and Q61H), we adapted an APEX2-dependent proximity labelling method **(Fig. 1A)**. To this end, we first generated tetracycline-inducible stable cell lines to minimize KRAS misfolding and mislocalization. Also, the same parental cell line was used as a control to generate a ‘benchmark’ proteome, enabling an unbiased comparison. Four different HEK293 FRT T-Rex cell lines (KRAS WT, G12D, G13D and Q61H) expressing WT KRAS- and all mutants- APEX2 fusion proteins in a stable fashion upon treatment with tetracycline **(Fig. 1B)** were generated. Furthermore, to examine the activity of APEX2 within the expressed constructs, streptavidin blot analysis and immunofluorescence were used to test its in-cell biotinylation ability. In the presence of tetracycline, phenol biotin and H_2_ O_2_, biotinylated proteins are detected as a wide band pattern, whereas just three endogenous biotinylated protein bands observed in all negative controls, indicating highly specific APEX2 labelling **(Fig. S1A)**.

**Figure 1:**
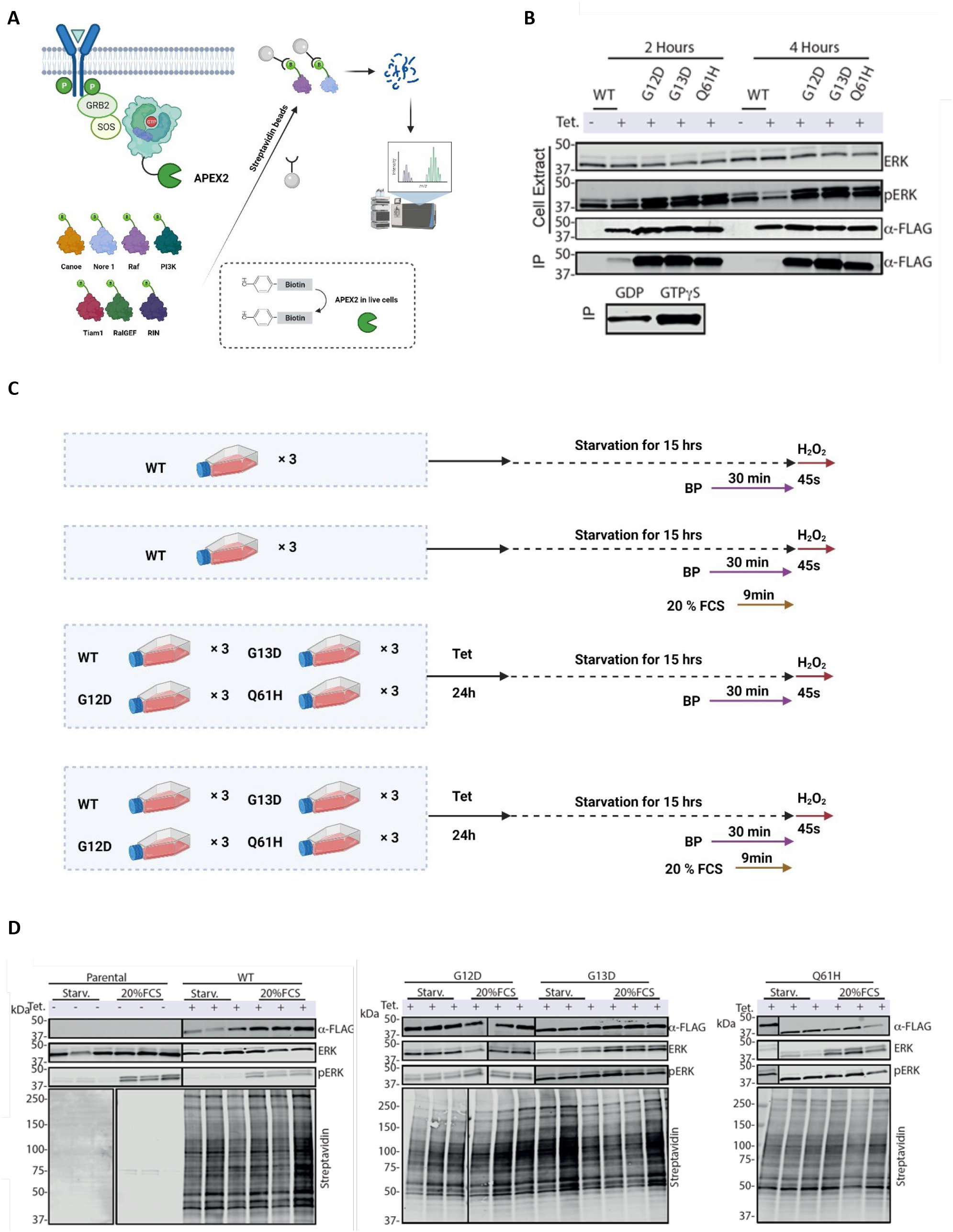
KRAS-APEX-2 wildtype, G12D, G13D and Q61H mutants express, localise and function similarly to endogenous KRAS. **(A)** Schematic representation of the approach used in this study to investigate the proximal proteome of K-Ras. First step is the generation of a fusion K-Ras APEX-2 protein with a flexible linker in between K-Ras and APEX-2. Second step, phenol biotin and H2O2 was added in the media to facilitate the biotinylation of proteins close proximal to the bait including effectors, regulators and interactors of K-Ras as well as proteins located in close proximity with K-Ras. The next step included the IP using streptavidin beads to capture biotinylated proteins. Finally, biotinylated proteins were digested, and mass spectrometry was used to identify proximal proteins. **(B)** Western blot analysis showing ERK, pERK, and a-FLAG in both cell extract and IP (immunoprecipitation) using GST-Raf1-RBD (Activate Ras Detection Kit) 2 and 4 hours upon starvation. As an IP positive control lysate was incubated with GTPγS and for IP negative control lysate was incubated with GDP. (**C**) Experiments were carried out in biological triplicates (n=3). KRAS has a long half-life the absent of tetracycline will decrease the proximal labelling of the new expressed KRAS. For the starvation condition cells were directly move for 30 minutes incubation with phenol biotin and subsequent incubation with H_2_ 0_2_ for 45 seconds. On the other hand, for the FCS induction condition cells were starver for 15 hours with a subsequent incubation with phenol biotin for 30 minutes. The last 10 minutes additional 20% FCS (including biotin phenol) was added to activate the cells. 45 seconds of H_2_ 0_2_ was used to initiate the biotinylating of the KRAS proximal proteins. **(D)** Western blot analysis showing ERK, pERK and a-FLAG in cell extract upon APEX-2 labelling under different environmental conditions including starvation and FCS induction. Additionally, western blot analysis indicating biotinylating proteins (Streptavidin, DyLight 488 Conjugated) upon streptavidin-immunoprecipitation. APEX-2 experiments were carried out in three independent biological experiments.

APEX-2 tagging may interfere with cellular localisation, signal transduction and GTP hydrolysis. To test this, the four cell lines were starved for 2 and 4 hours, respectively, and ERK/phospho-(p)-ERK activation was tested. As expected, upon starvation, pERK levels were reduced in WT KRAS, but not the mutants **(Fig. 1B)**. GTP-KRAS levels, measured using GST-Raf1-RBD enrichment, were decreased upon starvation only in the WT, but not in the constitutively active KRAS mutants, indicating that APEX- 2 does not interfere with these processes **(Fig. 1B)**. KRAS-APEX-2 fusion protein expression was further examined for its localisation. To this end, T-Rex tetracycline inducible lines expressing GFP-KRAS fusion protein and GFP only were generated (**Fig. S1B**). Microscopy analysis indicated the correct localisation of GFP-KRAS on the plasma membrane, whereas GFP only was located in the cytosol **(Fig. S1B)**.

Residual nuclear localisation was noted in less than 10% of the cells **(Fig. S1C)**. To complement this, subcellular fractionation was used to separate membranes from cytoplasmic fractions, confirming the localisation of KRAS-GFP and GFP only as observed by microscopy (data not shown). Also, the KRAS- APEX-2 WT and mutant fusion proteins were enriched in the membrane fraction **(Fig. S1D)**. We also examined the ability of KRAS-APEX2 fusion proteins to translocate upon starvation and FCS stimulation as their endogenous counterpart does. A Sucrose gradient fractionation experiment revealed similar translocation and localisation profiles for KRAS-APEX-2 fusion protein and endogenous KRAS upon 10 min FCS stimulation, 15 hours after initial starvation. Most importantly, APEX-2 labelling took place within the correct subcellular environment where endogenous KRAS is located **(Fig. S1E)**. Together, we conclude that KRAS-APEX-2 fusion proteins showed similar functional properties as compared to endogenous KRAS.

### APEX-2 labelling reveals differential molecular environments between WT and KRAS mutants

Proximity labelling experiments were conducted with APEX2-KRAS WT, G12D, G13D and Q61H expressing cell lines to compare their molecular vicinities. As controls, the APEX2-KRAS WT expressing cell line was analysed with and without tetracycline induction. We used a “beads only” background control as KRAS itself is expressed in different locations, thereby complicating analysis. In each case, we compared starvation and 10 min FCS induction. Tetracycline induction was performed overnight for a sufficient expression of the bait, followed by incubation in presence or absence of FCS for 15 hours **(Fig. 1C)**. Protein biotinylation was observed only in the presence of tetracycline, and the Raf/MEK/ERK pathway was not active during starvation of the WT, but not for the cells expressing oncogenic mutants (**Fig. 1D**).

Proximity labelled biotinylated material was isolated, digested and analysed by quantitative mass spectrometry. MS data were further processed with an interaction scoring algorithm SAINTexpress **(Teo et al., 2014)** to establish interaction probabilities (true proximity proteins) with KRAS baits under different conditions. A Saint Score above 0.8 with >=4- fold enrichment yielded 700-1000 KRAS proximal proteins per condition and a total of 1,373 proteins identified **(Fig. 2A,B)**.

**Figure 2:**
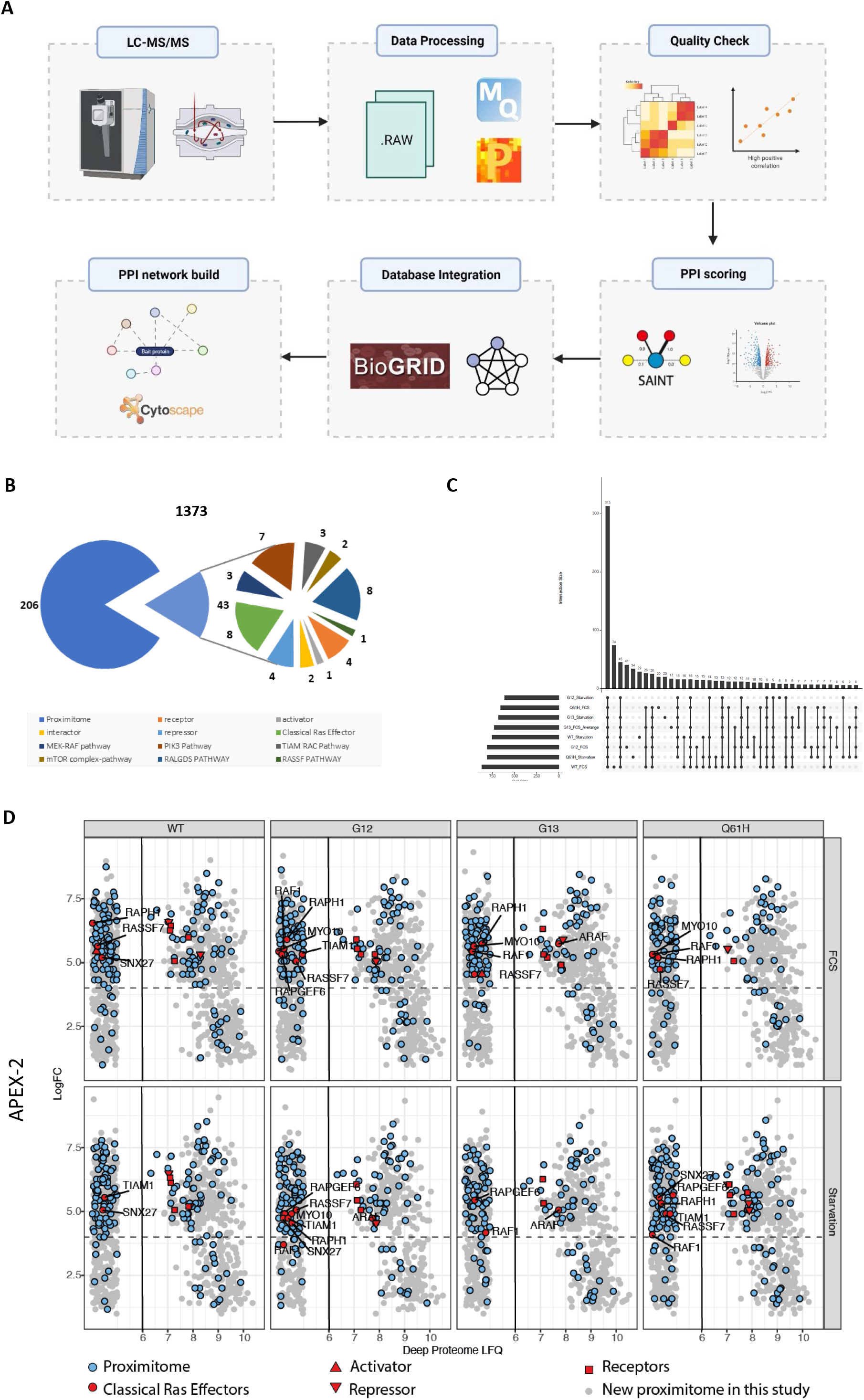
KRAS wildtype, G12D, G13D and Q61H proximal proteomes. **(A)** Mass spectrometry data analysis schematic representation. Data were initially analysed MaxQuant to identify proteins presence in our samples. Data were then processed with an interaction scoring algorithm SAINTexpress to provide a score on the probability of a true ‘interaction’ (true proximity protein) with our bait. Finally, true proximity proteins were decided to include any protein that has SAINT score above 0.8 and fold difference higher or equal to 4 as compared to the beads control. On the other hand, MaxQuant results were further analysed on Perseus and volcano blots were generated. **(B)** All Ras-proximal proteins identified in each APEX-2 KRAS proximitome. SAINTq probability ≥ 0.8; fold change ≥ 4 over beads control (1373 proteins). Several KRAS interactome families (Receptors, Repressors, Activators, Interactors, Classical Ras effectors, Proximitome, MEK-RAF pathway, PIK3 pathway, TIAM-RAC pathway, RALGDS pathway, RASSF pathway) were explored. **(C)** Histogram showing intersection of common protein groups between KRAS species and treatment. **(D)** KRAS proximal landscape (blue - previously known; grey – from this study) from the APEX-2 proximitome (left compartment) relative to the cellular proteome (right compartment). Previously known Ras effectors including activators, repressors and activators are indicated in red.

First, KRAS interacting partners were divided into several categories including receptors, activators, repressors, interactors and canonical RAS effectors proteins **(Kiel et al., 2021)**. Additionally, to “benchmark” our results, the BioGRID (https://thebiogrid.org/) database was used to retrieve the previously known KRAS ‘proximitome’ **(Adhikari and Counter, 2018; Bigenzahn et al., 2018)**. We detected proteins associated with KRAS signal transduction (Receptor Tyrosine Kinases (RTKs) including EGFR, MET and ERBB2IP), eight Ras effectors (RAF1, a-RAF1, TIAM1, RAPH1, RAPGEF2, RAPGEF2, SNX27 and MYO10). Also, several known KRAS interactors, activators and repressors were identified in our experiment including the E3 ligase adaptor protein LZTR1, DAG1 and GRB2 **(Fig. 2B)**. We also captured several proteins found to be part of downstream KRAS signalling, including proteins taking part in the networks of MEK-Raf (3 proteins), PIK3 (8) mTOR (2), TIAM RAC (3), RALGDS (8) and RASSF pathways (1) **(Fig.2 B).**

206 out of the 1,373 proximal proteins identified were shared between our study and previous BioID proximal studies **(Fig. 2B)**. 313 proteins were common between conditions and many hits unique to the different KRAS species **(Fig. 2C)**. APEX2 proximity labelling efficiency is reflected by the capture of low abundance proteins, not readily present in HEK293 deep proteome data sets **(Geiger et al., 2012)**. For instance, we captured several low abundant canonical KRAS ‘interactors’, such as RAF, TIAM1, SNX27, MARK2, demonstrating effective APEX2-based enrichment **(Fig. 2D)**.

Despite stringent threshold criteria, 1,373 identified proteins remained after the first pass, requiring further filtering approaches for the confident discovery of novel KRAS ‘proximitome’ candidates. A major hallmark revealed by our experiment, different from previous studies, was how the oncogenic mutations G12D, G13D and Q61H, located in the KRAS G domain, perturbed the KRAS molecular environment. For instance, in contrast to WT KRAS, RAF1 and ARAF were enriched with G12D, G13D and Q61H in both starvation and FCS stimulation **(Fig S3A-S3F)**, suggesting that the RAF-MEK-ERK pathway is highly activated in theses mutants, consistent with the ERK phosphorylation status **(Fig 1D)** and previous studies **(Haigis, 2017)**. It is noted that, not only the mutations but also the expression level can affect protein-protein interaction (PPI) networks. Therefore, the enhanced interaction between ARAF, RAF1 and KRAS mutants may also be caused by high expression. Furthermore, NF1 and SPRED2 are only enriched with the G13D and Q61H mutants, but not with G12D, suggesting that G13D and Q61H are associated with NF1/SPRED2 to a greater extent as compared to G12D, consistent with previous findings reporting that NF1 is not able to affect G12D KRAS **(Rabara et al., 2019)**. Notably, we found that LZTR1, an essential regulator for KRAS ubiquitination and degradation, was less abundant in all three mutant KRAS proximal proteomes, suggesting a decreased interacting affinity with mutant KRAS.

To further investigate the pathways that are selectively ascribed to or shared between the proximal proteins enriched in three different mutants, under resting or FCS stimulation conditions, we performed network enrichment analysis and visualization using Metascape **(Fig. 3A-3B)**. The network shows that processes such as Rho GTPase signalling, actin filament-based process, and cell morphogenesis were clustered and shared between resting and FCS stimulation conditions. Under the resting condition, pathway clusters such as cellular amino acids metabolism, actin filament-based process, are mostly shared between the G13D and Q61H mutants, indicating that the mutations potentially affect KRAS function via perturbations in proximal protein networks. In contrast, under FCS stimulation condition, most of the pathway clusters are shared among all three mutants, suggesting less mutant-specific effects.

**Figure 3:**
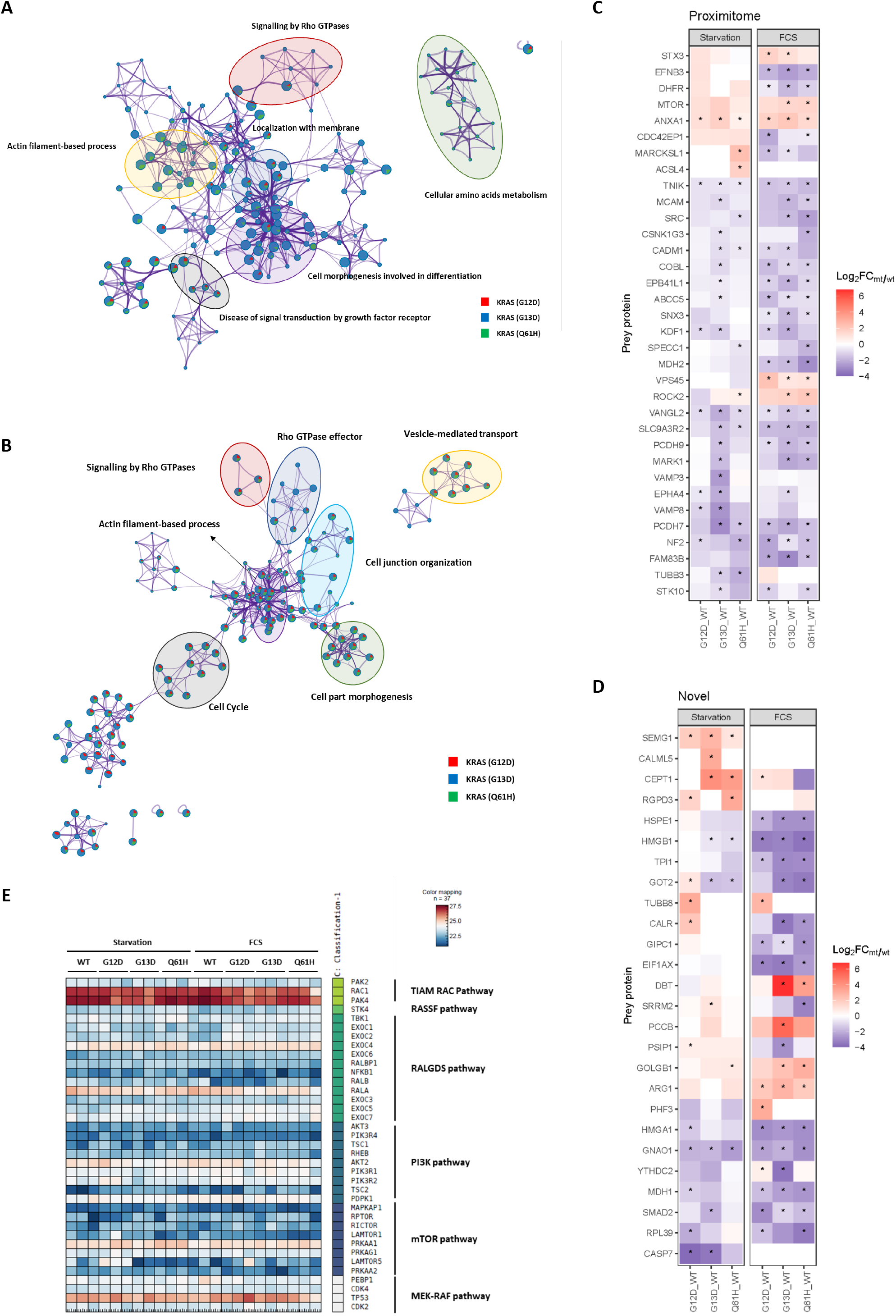
Enriched proteomes between KRAS wildtype and G12D, G13D and Q61H mutants. (**A-B**) Metascape enrichment network visualization showing the intra-cluster and inter-cluster similarities of enriched terms (up to ten terms per cluster), in starvation (**A**) and FCS stimulation state **(B)** separately, where nodes are represented by pie charts indicating their associations with three different mutants. The colour code for pie sector represents G12D (red), G13D (blue), and Q61H (green). Cluster labels were added manually. (**C**-**D**) Heatmap showing the log2(FC MUT/WT) of differentially enriched proximal proteins which were reported in previous proximitome studies (**C**) or proposed based on our study (**D**) Asterisk indicates that the hit meets the criteria of log2(mutant/WT FC) > 0.5 or <-0.5 and - log10 (p value) > 0.7. (**E**) Heatmap showing the relative abundance (log2 LFQ) of proximal proteins involved in KRAS related canonical pathways (such as MEK-RAF, mTOR, PI3K, RALGDS, RASSF and TIAM RAC pathway).

We also examinded changes between the ‘proximitome’ of the three KRAS mutants, especially in the newly identified proteins as well as the subset of hits that have previously reported in other proximity KRAS studies. We observed stronger association of mTOR, with all the three mutants as compared to the wild type, as expected. Interestingly, a stronger association with WT as compared to mutants was observed with NF2, a possible negative regulator of KRAS signalling (Cui et al.,2019). Several ‘proximitome’ hits seem to have a preference toward a specific KRAS mutants such as VPS45 (G12D FCS), ACSL4 (Q61H ST) VAMP3 (G13D ST) and ROCK2 (G13D and Q61H) **(Fig 3C)**. VAMP3 as part of SNARE complex was previously reported to regulate KRAS membrane localisation, identified to interact equally when we compare WT Vs G12D and Q61H **(Fig 3C)**. Nevertheless, during starvation, the G13D mutant seem to dramatically decrease its interaction with VAMP3 **(Fig 3C)**.

Any proteins that were not previously reported as a known KRAS proximitome or interactome were considered as novel KRAS proximal proteins. 26 were found to be enriched in one out of the three KRAS mutants as compared to wild type **(Fig 3D)**. Several of those proteins show a preference toward a specific KRAS mutant **(Fig 3D)**.

### LZTR1 differently affects wildtype versus KRAS mutants

This prompted us to perform a systematic search for KRAS associated partners with a different mutant associated abundance profile in our data set. Canonical and non-canonical interactors (including effectors, activators, repressors and receptors) were selected and visualised **(Fig 4A)**. Among the 105 previously reported physical interactors, 22 were identified here. Of these, LZTR1, ARAF, RAF1 and NF1 showed different abundance profiles between WT and mutant KRAS. Most notably, LZTR1 showed a weaker abundance associated with oncogenic G12D, G13D and Q61H KRAS mutants as compared to wildtype (Fig 4A). To test the preference of LZTR1 to be preferentially enriched with WT KRAS over the mutants, an APEX-2 labelling experiment was performed with Western blotting as a readout. Indeed, LZTR1 was detected in all samples except the beads control and enriched in the WT versus the mutants. ARAF was used as a control reflected by an enriched profile associated with KRAS mutants **(Fig. 4B)**.

**Figure 4:**
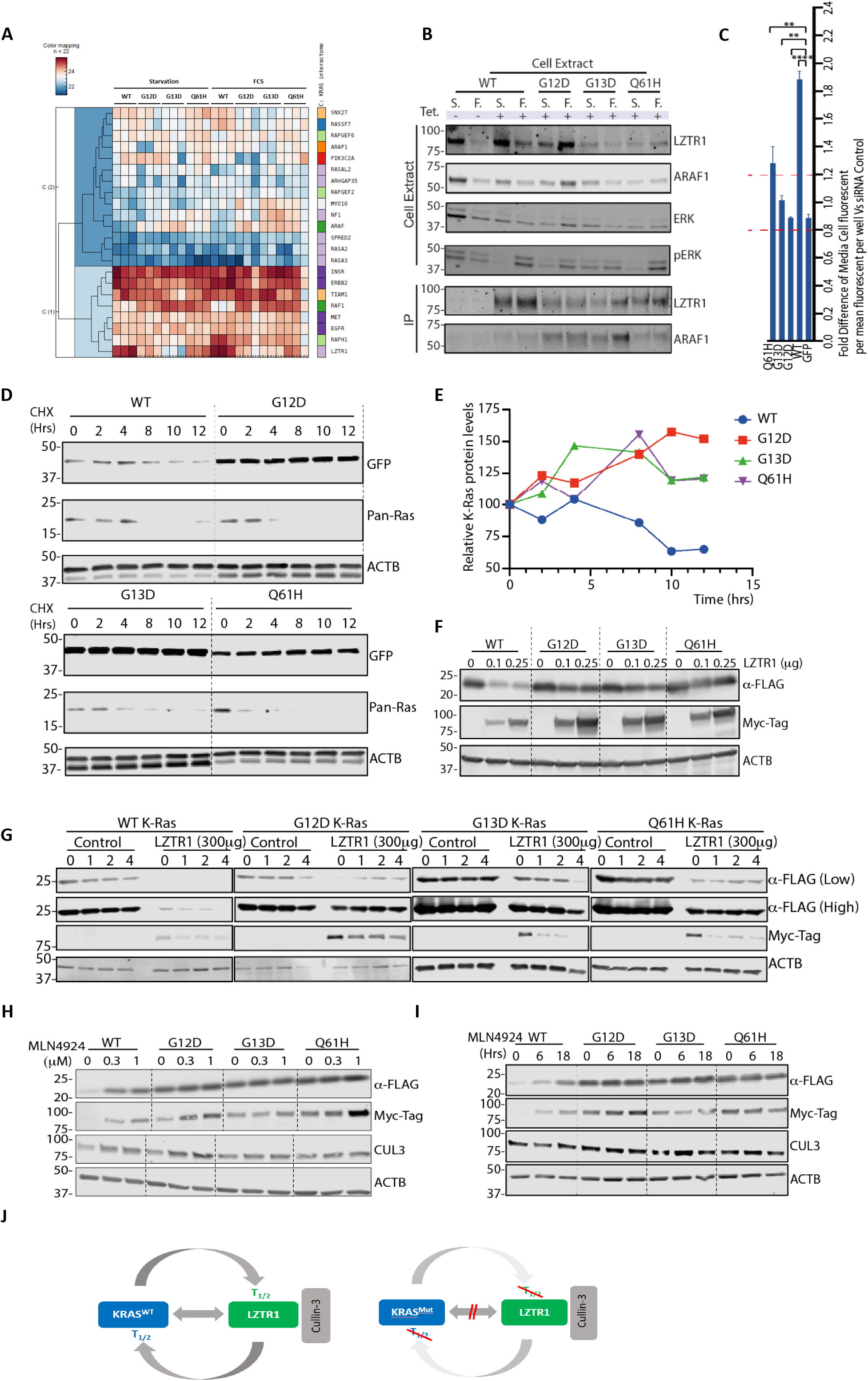
Interplay between KRAS and LZTR1 protein turnover, controlled by CUL3, is disrupted by KRAS mutations. **(A)** Heatmap showing relative abundance of known KRAS interactors in WT KRAS, G12D, G13D and Q61H KRAS mutant-APEX2 samples in either starvation or FCS stimulated condition. **(B)** APEX-2 labelling was performed in all different cell lines (WT, G12D, G13D and Q61H) under starvation and FCS induction. Western blot analysis showing ERK, pERK, ARAF and LZTR1 in cell extract as well as elution upon streptavidin-immunoprecipitation. **(C)** Cells were seeded in 96 well plates and treated with tetracycline for 24 hours. Next day cells were transfected with LZTR1 siRNA or Control siRNA. Cells were imaged after 72 hours using the Opera Phenix microscope. The images were analysed in Columbus Image Data Storage and Analysis System. The media fluorescent per well per mean per cell was determined. Finally, a fold difference between the control and the siRNA LZTR1 was measured. **(D)** Stable HEK293 FRT T-REX cell lines of WT, G12D, G13D and Q61H were treated with tetracycline for 24 hours. The next day, cells were treated with 50 μg/mL cycloheximide (CHX). At the indicated time points after CHX-treatment cells were harvested. Western blot analysis showing GFP, Pan-Ras and ACTB in cell extract was performed. **(E)** Band densities from Figure 4D were analyzed using ImageStudio, and expression was normalized to ACTB protein levels to determine KRAS half-life. **(F)** The influence of LZTR1 overexpression on WT K-Ras as well as G12D, G13D and Q61H oncogenic mutants. HEK293 cells were seeded in 6 well plates and transfected with 1µg Flag-K-Ras plasmids as well 0, 0.1 or 0,25 µg Myc-LZTR1 plasmid concentration. Cells were cultured for a total of 48 hours before protein levels were evaluated using Western blot. **(G)** HEK293T cells were transfected with 1 µg Flag-RAS and 0.2 µg Myc-LZTR1 plasmid concentration. Cells were incubated for 48 hours, then cells were treated with 50 μg/mL CHX. At the indicated time points after CHX-treatment cells were harvested. Western blot analysis showing FLAG (RAS), MYC (LZTR1) and ACTB in cell extract was performed. **(H)** HEK293T cells were transfected with 1 µg Flag-RAS and 0.2µg Myc-LZTR1 plasmid concentration and let to grow for 48 hours. Cells were then treated for 24 hours with MLN4924 in different concentrations (0, 0.3 1 µM). Cell extract was then analysed on western blot and FLAG (RAS), MYC (LZTR1), CUL3 and ACTB was monitored. **(I)** HEK293T cells were transfected with 1 µg Flag-RAS and 0.2 µg Myc-LZTR1 plasmid concentration and let to grow for 48 hours. Cells were then treated for indicated time with MLN4924 (0.3 µM). Cell extract was then analysed on western blot and FLAG (RAS), MYC (LZTR1), CUL3 and ACTB was monitored.

### LZTR1 controlled KRAS protein turnover is altered by oncogenic KRAS mutants

The enrichment of LZTR1t in the proximal proteome of WT KRAS related to the mutants might be linked to differential protein turnover. To test this, siRNA knockdown of LZTR1 was performed in HEK293 cells expressing WT KRAS-GFP, G12D-GFP, G13D-GFP and Q61H-GFP fusion proteins. As a control, a GFP only HEK293 FRT cell line was generated. The GFP fluorescent signal was monitored in the different stable cell lines, and the GFP cell line was used to assess the global effect of LZTR1 on protein expression. Knockdown of LZTR1 stabilised WT KRAS GFP expression but not GFP only **(Fig.4C, S5A),** consistent with its role as an adaptor protein to promote ubiquitination of KRAS, resulting its degradation as reported previously **(Abe et al., 2020)**. Notably, KRAS stability was strongly diminished in the G12D, G13D KRAS and slightly decreased in the Q61H KRAS mutants **(Fig.4C, S5A)**. This suggests that the half-life of oncogenic KRAS mutants might be increased compared to the WT counterpart. To further explore this, KRAS-GFP cell lines expressing WT and oncogenic KRAS mutants (G12D, G13D and Q61H) were treated with cycloheximide (CHX) for different times. We observed a shorter half-life of WT KRAS as compared to the oncogenic mutants **(Fig. 4D, E)**. WT KRAS was strongly depleted after 10 hours of CHX treatment, and the oncogenic mutants more stable **(Fig. 4D, E)**. To further explore whether LZTR1 differently affects KRAS oncogenic mutants, Myc-LZTR1 was co-expressed with 3Flag- KRAS WT, G12D G13D or Q61H. Elevated LZTR1 levels yielded substantially decreased WT KRAS protein levels, whereas the three oncogenic KRAS mutants show little to no protein downregulation. Most notably, LZTR1 protein levels were highly upregulated in the presence of oncogenic KRAS, suggesting a possible interdependence between KRAS and LZTR1 protein levels and turnover **(Fig. 4E)**. To get further insight, we tested how KRAS mutations affect protein stability in the presence of LZTR1 using a cycloheximide (CHX)-chase assay. At the start of the CHX-chase assay, WT KRAS protein levels were much lower as compared to any of the three mutants. In cells co-expressing Flag- KRAS WT and Myc- LZTR1, KRAS protein levels were reduced by half after 12 hours of CHX treatment, whereas the three mutant protein levels were unchanged. Remarkably, LZTR1 levels were also differentially affected in the presence of WT versus mutant KRAS **(Fig. 4G)**. Taken together, these results suggest that KRAS degradation, occurring in an LZTR1 dependent manner, is altered by oncogenic mutations. In turn, LZTR1 levels are also affected in the presence of WT, but not KRAS mutants.

### WT KRAS and LZTR1/CUL3 regulatory circuit disrupted by oncogenic KRAS mutants

LZTR1 was described as a substrate adaptor of CUL3-based ubiquitin E3 ligase **(Bigenzahn et al., 2018; Hollstein and Cichowski, 2013).** Therefore, we decided to examine whether the inhibition of NEDD8- activating enzyme (NAE) by inactivation of Cullin-RING E3 ubiquitin Ligases (CRLs) via blocking Cullin neddylation has any effect on KRAS and LZTR1 protein stability. MLN9424 is a newly discovered small molecular inhibitor of NEDD8-activating enzyme (NAE) and it was used as an inhibitor of CRLs **(Milhollen et al., 2010)**. HEK293 cells were initially co-transfected with LZTR1 and WT or KRAS mutants, followed by treatment with different concentration of MLN9424 for 24 hours. Co-transfection of LZTR1 and WT KRAS reduced KRAS protein levels, which were restored in the presence of MLN9424 **(Fig. 4H)**. This was not observed with the oncogenic KRAS G12D and G13D mutants, perhaps to some extent with Q61H in the presence of high concentrations of MLN9424 (1μM). Notably, LZTR1 protein levels were upregulated with MLN9424, predominantly in presence of WT KRAS. LZTR1 and WT KRAS protein levels showed an interrelated abundance profile not observed with the mutants, perhaps with the exception of KRAS G12D to a minor extent **(Fig. 4H)**.

To study the underlying dynamics of this effect, MLN9424 inhibitor experiments were performed in a time-dependent fashion. HEK293 cells were co-transfected with Myc-LZTR1 and Flag-KRAS. Cells were then incubated with 0.3 μM MLN9424 for 0, 6 and 18 hours, respectively. WT KRAS protein levels were gradually increased, whereas the oncogenic KRAS mutant protein levels remained at a similar level **(Fig. 4I)**. LZTR1 protein levels followed a comparable trend. Once more, only in the presence of the WT KRAS, LZTR1 protein levels were notably increased. A small exception was G12D KRAS, were a minor increased in LZTR1 levels was observed **(Fig. 4I)**. Since Cullin3 was previously shown to regulate (WT) KRAS protein levels via LZTR1 **(Abe et al., 2020),** we knocked down Cullin 3 to investigate how LZTR1 protein levels behaved in the presence of CHX. Removal of Cullin3 led to an increase of LZTR1’s protein half-life, suggesting that Cullin 3 regulates its own co-factor (LZTR1) in a substrate (KRAS)- dependent manner **(Fig. S5B)** To further examine how KRAS protein levels affect LZTR1 protein stability, HEK293 cells were co-transfected with LZTR1 and KRAS, where either KRAS **(Fig. S5C)** or LZTR1 plasmid concentration were kept constant **(Fig. S5D)**. In both cases, we observed that LZTR1 protein levels were stabilised in a dose-dependent manner similar to KRAS, indicating a direct crosstalk between KRAS and LZTR1. This appeared to be Cullin3 dependent, because when Cullin 3 was knocked down, Flag-KRAS and Myc-LZTR1 co-expression did not lead to LZTR1 accumulation, whereas Flag-KRAS protein levels were stabilised **(Fig. S5E)**. We conclude that the regulation of KRAS and LZTR1 turnover are linked, possibly involving Cullin Ring Ubiquitin Ligases (CRLs), in particular Cullin 3 (**Fig. 4J**).

## DISCUSSION

Oncogenic KRAS G12D G13D and Q61H are amongst the most common point mutations found in human cancer **(Cook et al., 2021; Liu et al., 2019)**. KRAS mutations appear in different cell types with allele-specific profiles, suggesting different underlying mechanisms behind driving cell transformation. For instance, G12D KRAS mutation is appearing in over 90% of pancreatic ductal adenocarcinoma (PDAC) **(Wong et al., 2016),** G13D is most frequent in colon adenocarcinoma **(Taieb et al., 2017)** and Q61H in lung adenocarcinoma **(Kunimasa et al., 2020).** KRAS-dependent oncogenesis is depended on at least four factors contributing to different KRAS mutations being associated with specific cancer types **(Prior et al., 2012)**. First, expression levels as well as GTP/GDP-KRAS ratios contribute to cell transformation. Active KRAS dynamics varies considerably between different oncogenic mutations **(Killoran and Smith, 2019)** and maybe influenced by alterations in GEF, GAF and effector interactions **(Kiel et al., 2021).** Finally, the genetic, epigenetic and proteomic environment of different cell types differential impact RAS-dependent oncogenesis.

Here, we systematically investigated how these oncogenic driver mutations of KRAS affect their proximal molecular environment. The use of an isogenic cell line as control reduced interference due to different cellular backgrounds. Our approach relied on an over-expression system, and it is known to potentially provoke cell transformation **(Ito et al., 2021).** However, we have carefully controlled this with an optimised starvation protocol to mimic a “baseline state”. Of note, we cannot exclude the possibly of missing crucial KRAS associations in our experimental system due to intrinsic limitations of over-expression systems. An APEX-2 proximity labelling was used as it occurs at a timescale of seconds, which is advantageous for the capture of transient interactions as well as fast molecular changes as compared to BioID-based approaches **(Trinkle-Mulcahy, 2019)**. Labelling conditions were optimised and levels of protein biotinylation assessed in whole cell lysates as well as streptavidin eluate material to control for consistency (**Fig 1D**). Previous proteomics-based studies exploring the KRAS “proximitome”, based on BioID, revealed many shared proteins between WT & mutant KRAS (mainly G12V, S17N, C185S), predominantly involved in protein expression and trafficking **(Kiel et al., 2021; Li et al., 2012; Matallanas et al., 2003)**. Additionally, there appears to be a large overlap between GTP/GDP KRAS interactors (proteins in closed proximity), indicating a more gradual KRAS activation transition rather than being an on/off switch. Very few specific interactors were reported, such as the KRAS G12V selective interaction with phosphatidylinositol-4-phosphate 5-kinase type 1 (PIP5K1), mediating PI3K and ERK signalling **(Semenas et al., 2014)**.

Our approach offers a deeper insight as APEX2 may capture more concise molecular snapshots, and we systematically compared the molecular environment of WT KRAS compared to G12D, G13D and Q61H mutants, respectively, under resting and acute FCS stimulation conditions. Residues G12, G13 and Q61 are in the KRAS G domain, altering nucleotide binding and exchange and are expected to alter KRAS interacting networks. To further examine the difference in the downstream network, proteins involved in the known KRAS signalling pathways, the RAF-MEK, mTOR, PI3K, RALGDS, RASSF and TIAM-RAC networks were selected and visualised **(Fig 3G)**. In total, 37 downstream signalling proteins were identified among all the samples, and we did not observe significant differences between WT KRAS and all the mutants. Gene ontology-cellular component analysis for WT and the mutants showed enrichment of focal adhesion and plasma membrane, in line with its subcellular localization **(Fig 3H)** and consistent with previous studies **(Kiel et al., 2021).**

The absence of some canonical Ras interactors, such as b-RAF, H-Ras/N-Ras and SOS, could be explained by the lack of accessible lysine residues for the biotinylation reaction to take place or perhaps the low expression level of some of those proteins in HEK293 cells, as it is the case for b-Raf **(Geiger et al., 2012)**. A-RAF in turn seems to be strongly enriched with KRAS G13D as compared to G12D and Q61H. On the other hand, NF1 and SPRED2 seem to be enriched to a greater extent with the G13D and Q61H mutants as compared to G12D, in line with previous findings that the G12D KRAS signalling pathway is not affected by NF1 **(Cheng et al., 2021)**.

As shown previously **(Bigenzahn et al., 2018; Kovalski et al., 2019)**, LZTR1 was detected in the micro- environment of both WT and selected KRAS mutants. Most notably, we found that LZTR1 levels were strongly reduced in proximity of the mutants, especially G12D, G13D and to a lesser extent with Q61H (**Fig 4A)**. LZTR1’s effect on ubiquitination and turnover of WT KRAS and some mutants has been examined (Abe et al., 2020). We extend this observation to the KRAS mutants G12D, G13D and Q61H, whose turnover was less dependent on LZTR1 (**Fig 4D, E**). Importantly, we discovered a reciprocal effect on LZTR1 protein turnover itself that was most prevalent in the presence of WT KRAS (**Fig 4B, D-I**) and CUL3-dependent (**Fig S5C, F**), indicating a direct (WT)-KRAS-LZTR1 regulatory circuit that appears to be disrupted by KRAS mutants (**Fig 4J**). This further demonstrates the interplay between LZTR1, a CUL3 ubiquitin E3 ligase adapter for KRAS. It will be interesting to explore whether LZTR1 mutations, prevalent in Rasopathies such as glioblastoma & dominant Nooan syndrome **(Motta et al., 2019; Steklov et al., 2018)**, may also affect this molecular regulatory circuit with potential consequences for malignant cell transformation.

## Acknowledgements

We thank members of the Kessler group and ITEN teams for helpful comments and insightful discussions. This work was supported by Pfizer. Work in the BMK lab was funded by the Chinese Academy of Medical Sciences (CAMS) Innovation Fund for Medical Science (CIFMS), China (grant number: 2018-I2M-2-002) and an EPSRC grant EP/N034295/1.

## Author contributions

AD and BMK directed this study. AD, ZL and GV performed biochemical and biological experiments. SH and RF performed mass spectrometry analysis. FL, GV, PD, ASD, APF and AD analysed and interpreted the data. AD, ZL and BMK wrote the manuscript. All co-authors made comments on the manuscript.

## FIGURE LEGENDS

**Figure S1:**
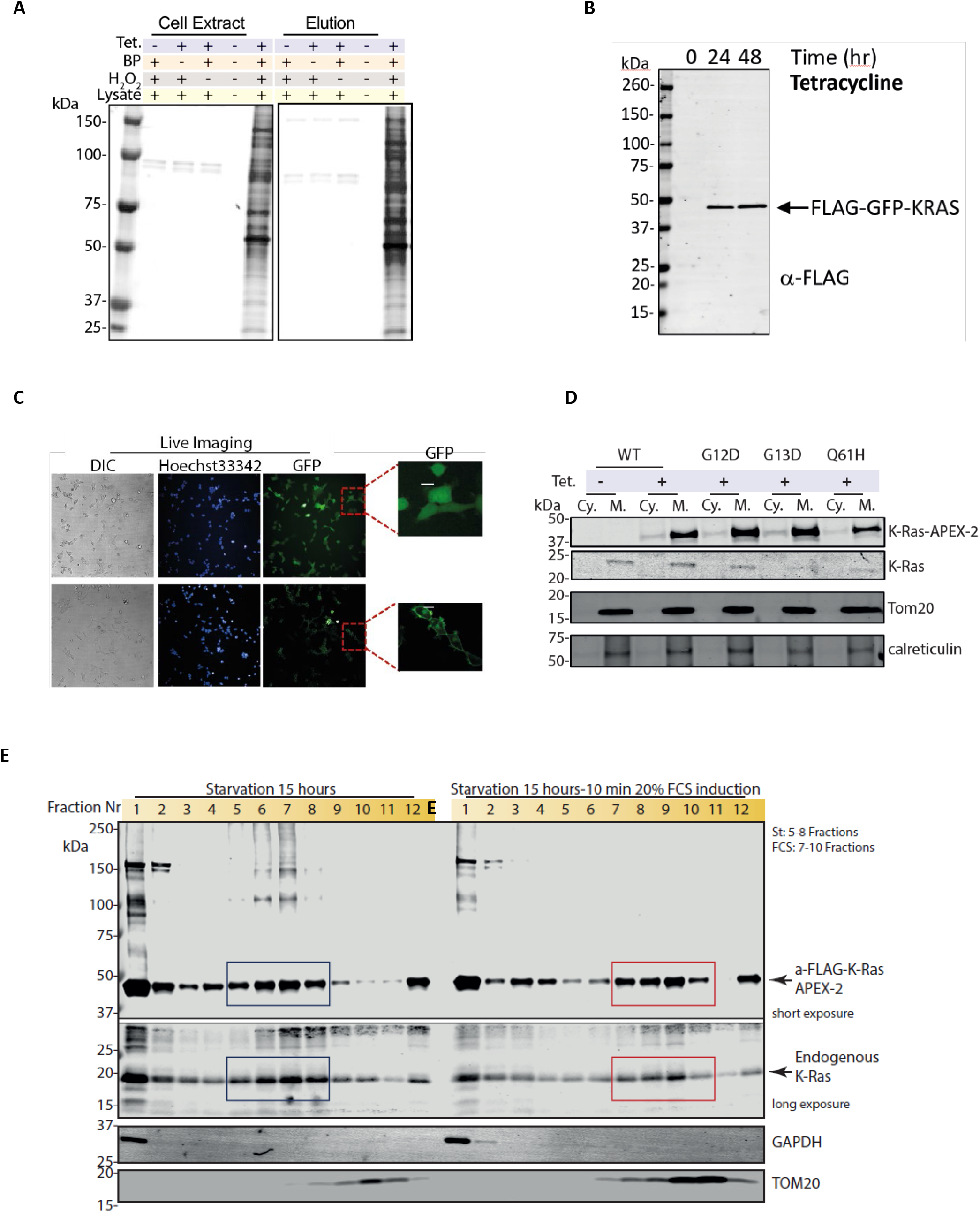
Endogenous, APEX2-KRAS and GFP-KRAS association with membrane components. **(A)** Western blot indicates the biotinylated proteins in the presence or absence of different c reagents including tetracycline, phenol-biotin and H_2_ O_2_. K-Ras APEX-2 stable cell line was treated with tetracycline for 24 hours where is indicated. Then, cells were treated with phenol-biotin and/ or H_2_ O_2_ (where is indicated) and cells were lysed, and proteins were separated in an SDS Gel. Streptavidin, DyLight 488 Conjugated antibody were used to visualise the biotinylating proteins. * Represents biotinylated background proteins. **(B)** Western blot analysis showing a-FLAG in cell extract upon 24 hours tetracycline treatment. **(C)** Imaging of live cells using Opera Phenix™ and processed in Columbus Image Analysis System. Panels 1183 Hoechst 33342 for staining DNA (blue); the green fluorescence signal from GFP (Green), Differential Interference Contrast (DIC). **(D)** Western blot analysis showing K-Ras GFP, GAPDH and GFP expression upon digitonin cytoplasmic-membrane fractionation of Stable K-Ras GFP or GFP only FRT T-Rex HEK293 cell line. Cells were incubated with tetracycline for 24 hours. **(E)** Western blot analysis showing a-FLAG, endogenous KRAS, TOM20 and Calreticulin expression upon digitonin cytoplasmic-membrane fractionation of stable KRAS APEX-2 WT, G12D, G13D and Q61H FRT T-Rex HEK293 cell lines. Cells were incubated with or without tetracycline for 24 hours. **(F)** Western blot analysis showing a-FLAG, GAPDH, TOM20 and endogenous KRAS in sucrose gradient fractions upon 15 hours of starvation or 15 hours of starvation and a subsequent 10-minute treatment with 20% FCS.

**Figure S2:**
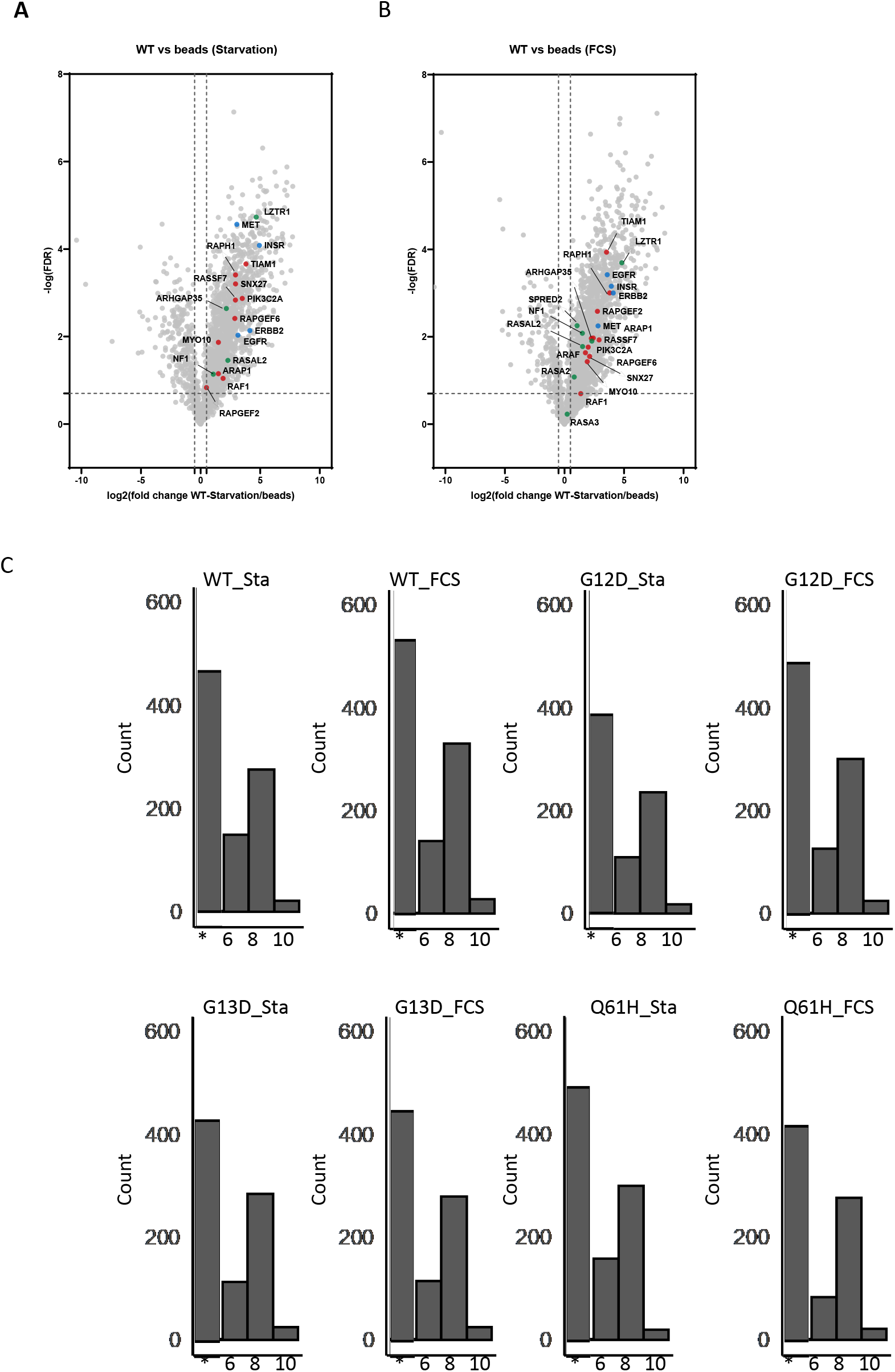
Enriched proteomes between KRAS wildtype exposed to starvation or acute FCS stimulation. (**A-B**) Volcano plotting of fold changes and p values derived from t-test statistic for proximal proteins identified in WT KRAS-APEX2 in either starvation (**A**) or FCS treated condition (**B**), in which the beads were used as a control. Known KRAS effectors (red), repressors (green) and receptors (blue) are coloured and labelled. (**C**) Histogram of proteins with differential protein abundances, reflecting enrichment per KRAS species.

**Figure S3:**
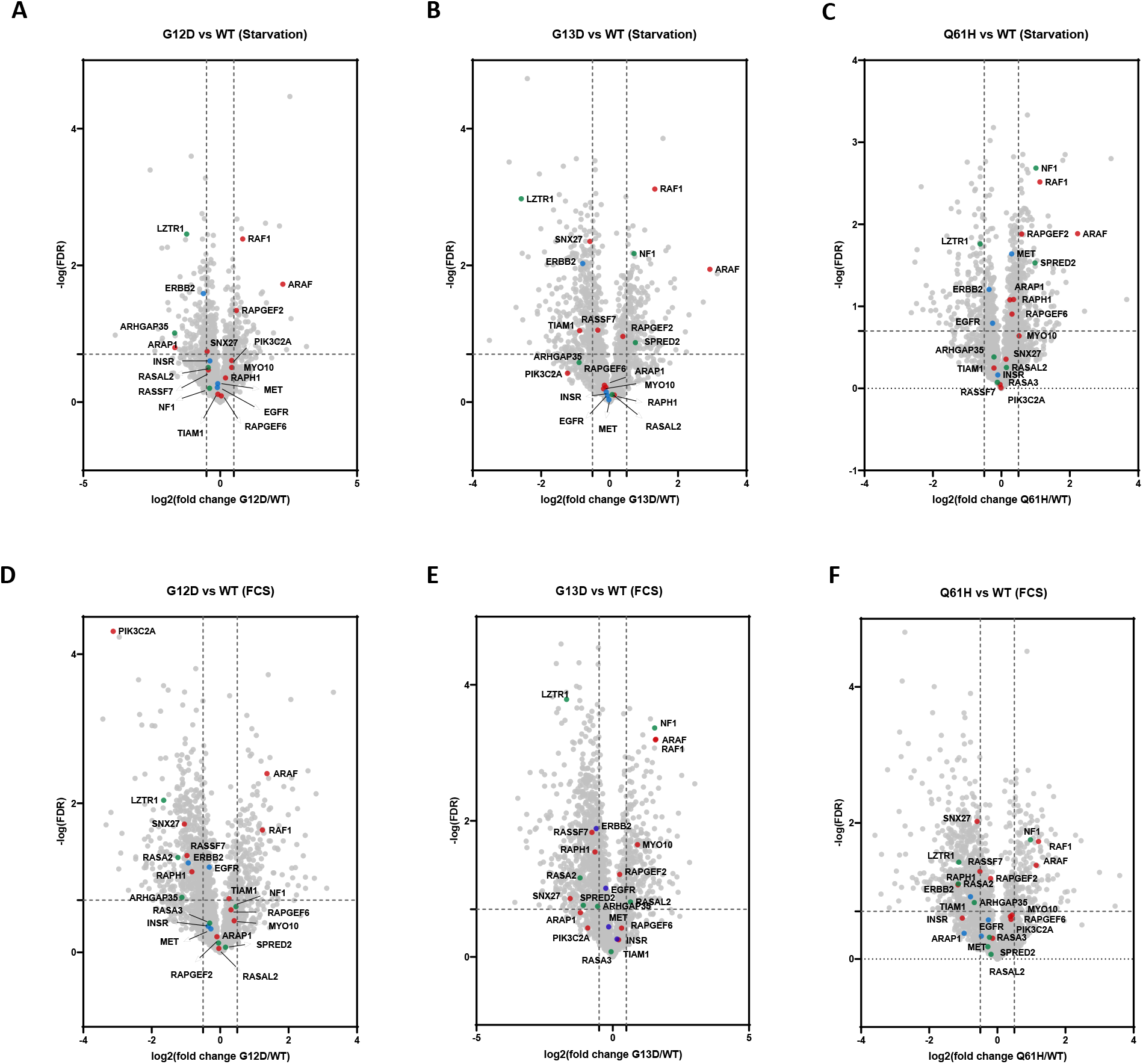
Enriched proteomes between KRAS wildtype and G12D, G13D and Q61H mutants. (A-F) Volcano plotting of fold changes and p values derived from t-test statistic for proximal proteins identified in G12D, G13D, and Q61H KRAS mutant in either starvation or FCS treated condition, in which WT-KRAS was used as a control. Known KRAS effectors (red), repressors (green) and receptors (blue) are coloured and labelled.

**Figure S4:**
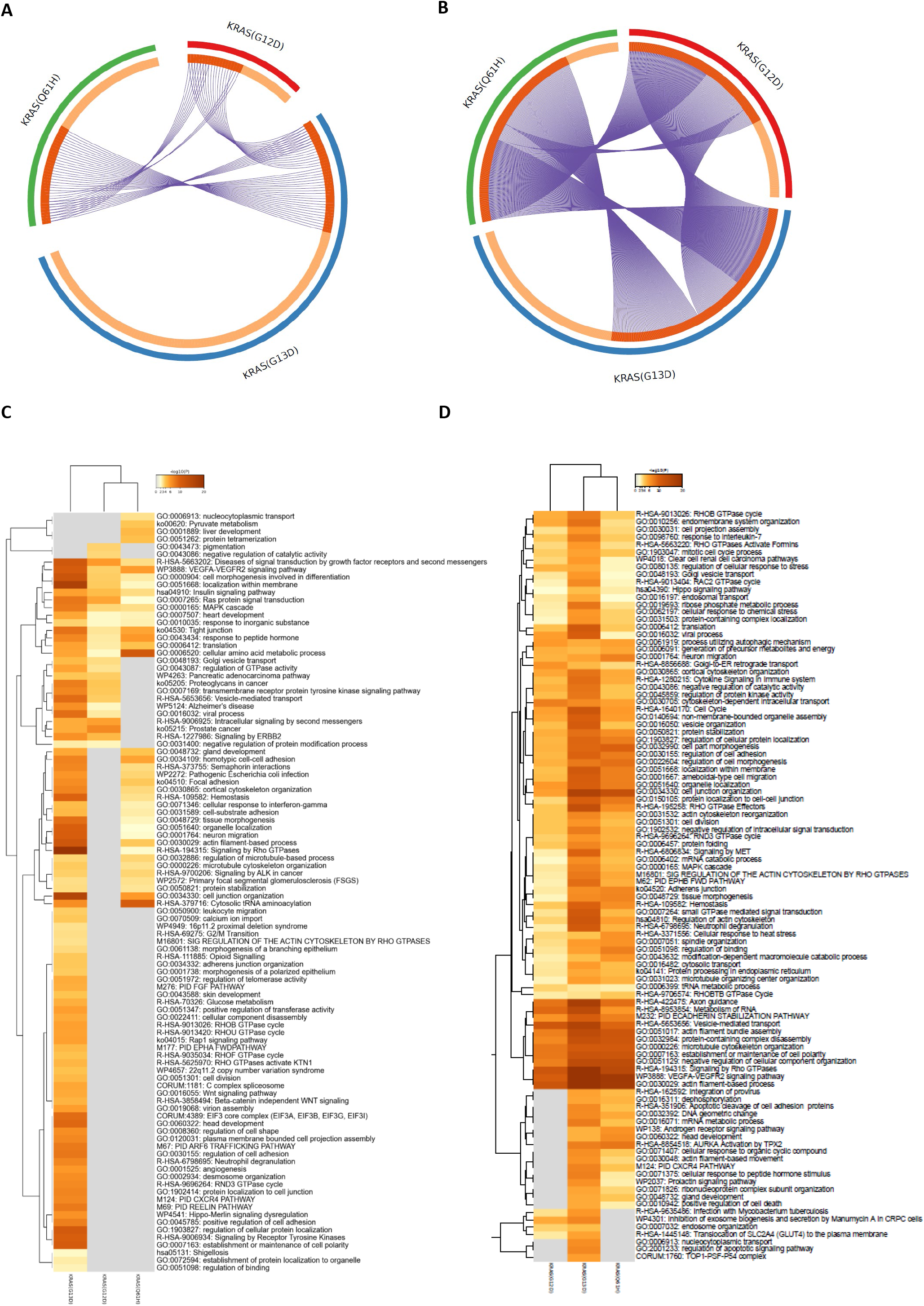
KRAS wildtype, G12D, G13D and Q61H protein interaction networks. (**A-B**) Circos plot showing the overlap of the proximal proteins identified from three mutants in starvation (**A**) and FCS stimulation state (**B**). (**C**-**D**) Heatmap showing the top enrichment clusters with colour scale representing statistical significance, which was performed using Metascape. Gray colour indicates a lack of significance.

**Figure S5:**
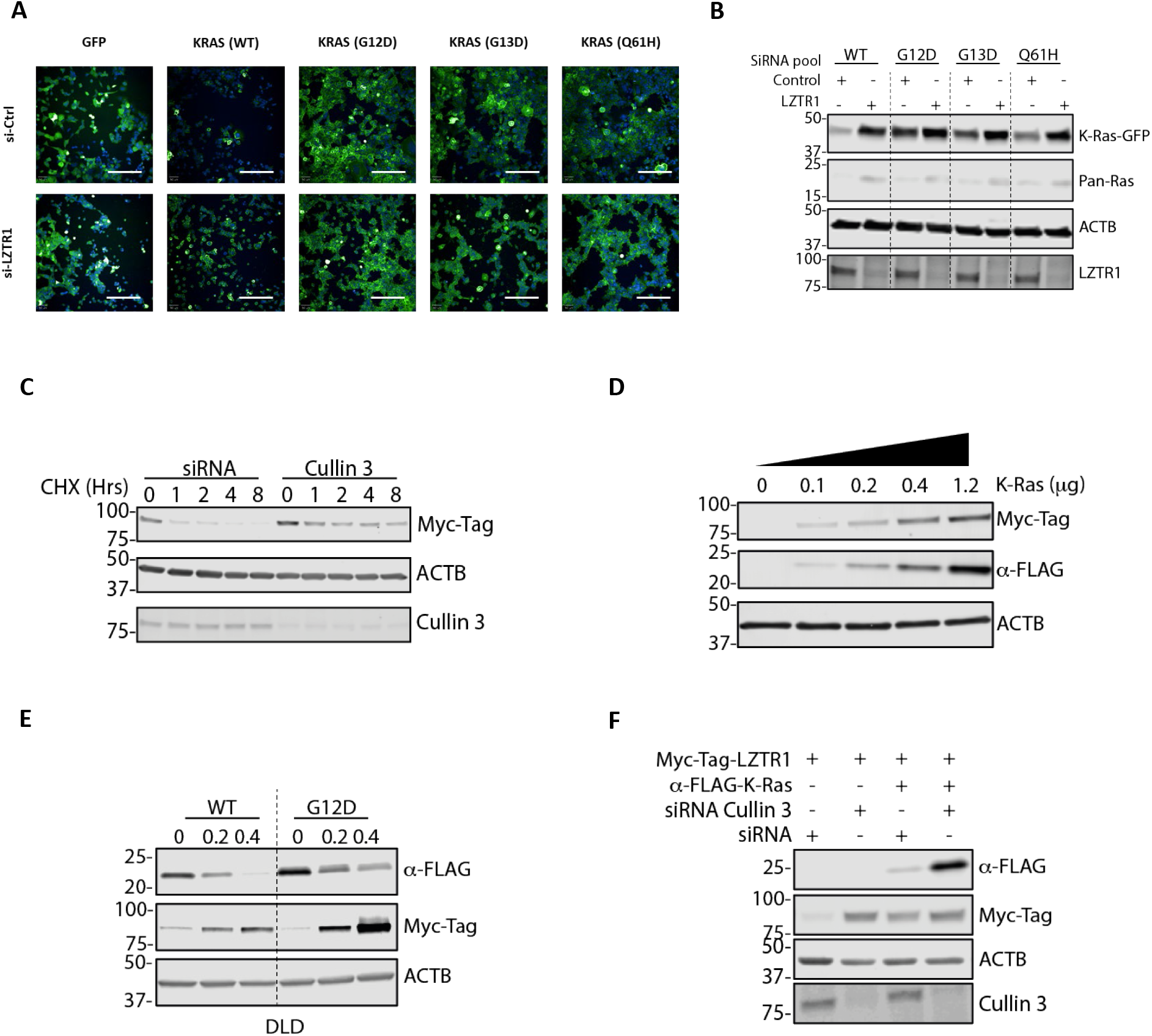
Interdependence of LZTR1 and KRAS protein stabilities controlled by CUL3. **(A)** Microscopy based analysis of GFP-KRAS in the presence and absence of LZTR1. (**B**) KRAS and Pan-Ras protein stability in the presence and absence of LZTR1 assessed by immunoblotting. (**C**) Cullin 3 (CUL3) knockdown stabilises LZTR1. (**D**) LZTR1 (MYC) and KRAS (FLAG) levels are correlated at the protein level. (**E**) KRAS wildtype and G12D (both FLAG tagged) differentially affect LZTR1 (MYC-tag) protein levels. (**F**) KRAS (FLAG) and LZTR1 (MYC) protein levels are dependent on the presence of Cullin 3 (CUL3).

